# Genomic modeling as an approach to identify surrogates for use in experimental validation of SARS-CoV-2 and HuNoVs inactivation by UV-C treatment

**DOI:** 10.1101/2020.06.14.151290

**Authors:** Brahmaiah Pendyala, Ankit Patras, Doris D’Souza

## Abstract

Severe Acute Respiratory Syndrome coronavirus-2 (SARS-CoV-2) is responsible for the COVID-19 pandemic that continues to pose significant public health concerns. While research to deliver vaccines and antivirals are being pursued, various effective technologies to control its environmental spread are also being targeted. Ultraviolet light (UV-C) technologies are effective against a broad spectrum of microorganisms when used even on large surface areas. In this study, we developed a pyrimidine dinucleotide frequency based genomic model to predict the sensitivity of select enveloped and non-enveloped viruses to UV-C treatments in order to identify potential SARS-CoV-2 and human noroviruses surrogates. The results revealed that this model was best fitted using linear regression with r^2^=0.90. The predicted UV-C sensitivity (D_90_ - dose for 90% inactivation) for SARS-CoV-2 and MERS-CoV was found to be 21 and 28 J/m^2^, respectively (with an estimated 18 J/m^2^ as published for SARS-CoV-1), suggesting that coronaviruses are highly sensitive to UV-C light compared to other ssRNA viruses used in this modeling study. Murine hepatitis virus (MHV) A59 strain with a D_90_ of 21 J/m^2^ close to that of SARS-CoV-2 was identified as a suitable surrogate to validate SARS-CoV-2 inactivation by UV-C treatment. Furthermore, the non-enveloped human noroviruses (HuNoVs), had predicted D_90_ values of 69.1, 89 and 77.6 J/m^2^ for genogroups GI, GII and GIV, respectively. Murine norovirus (MNV-1) of GV with a D_90_ = 100 J/m^2^ was identified as a potential conservative surrogate for UV-C inactivation of these HuNoVs. This study provides useful insights for the identification of potential nonpathogenic surrogates to understand inactivation kinetics and their use in experimental validation of UV-C disinfection systems. This approach can be used to narrow the number of surrogates used in testing UV-C inactivation of other human and animal ssRNA viral pathogens for experimental validation that can save cost, labor and time.

## 1 Introduction

Coronaviruses belong to the family of *Coronaviridae*, comprising of 26 to 30 kb, positive-sense, single-stranded RNA, in an enveloped capsid (Woo et al., 2010). Coronaviruses can cause severe infectious diseases in human and vertebrates, being fatal in some cases. Severe acute respiratory syndrome (SARS) coronavirus (SARS-CoV-1), a β-coronavirus emerged in Guangdong, southern China, in November, 2002 (Guan et al., 2003), and the Middle East respiratory syndrome (MERS) coronavirus (MERS-CoV), was first detected in Saudi Arabia in 2012 (Alagaili et al., 2014). Since late December 2019, a novel β-coronavirus (2019-nCoV or SARS-CoV-2) has been responsible for the pandemic coronavirus disease (COVID-19) with > 7.2 million confirmed cases throughout the world, and a fatality rate of approximately 5.7 % as of 11 June, 2020 (WHO, 2020a). This 2019-nCoV is thought to have originated from a seafood market of Wuhan city, Hubei province, China, and has spread rapidly to other provinces of China and other countries (Zhu et al., 2020).

According to current evidence documented by the World Health Organization (WHO), SARS-CoV-2 virus (2019-nCoV) is transmitted between humans through respiratory droplets and contact (person-to-person, fomites, etc.) routes (WHO, 2020b). van Doremalen et al. (2020) reported that SARS-CoV-2 remained viable in aerosols throughout the 3 h duration of the experiment and more stable on plastic and stainless steel than on copper and cardboard, and virus was detected up to 72 hours after the application to these surfaces at 21-23°C and 40% relative humidity. Given the ability of these viruses to survive in the environment, appropriate treatment strategies are needed to inactivate SARS-CoV-2. As per WHO recommendations, SARS-CoV-2 may be inactivated using chemical disinfectants. As of 07 April, 2020, the United States Environmental Protection Agency (USEPA) has announced a list of 428 registered chemical disinfectants that have been approved for use against SARS-CoV-2 (USEPA, 2020). However, chemical disinfection requires intense labor and product to treat large surface areas. As an alternative, ultraviolet light (UV) technology (with germicidal UV-C at wavelengths from 100 nm to 280 nm) can be an effective approach to inactivate SARS-CoV-2 on large surface areas and in the air (regardless of humidity levels) with less labor. UV inactivates a broad spectrum of microorganisms by damaging the DNA or RNA and thereby prevents and/or alters cellular functions and replication (Patras et al., 2020). UV-C inactivation of various microorganisms such as pathogenic bacteria, spores, protozoa, algae and viruses has been reported (Malayeri et al., 2016; Bhullar et al., 2019; Gopisetty et al., 2019; Pendyala et al., 2019, 2020; Patras et al., 2020). As of date, information on the UV susceptibility of SARS-CoV-2 is unknown.

Human noroviruses (HuNoVs) cause > 80 % of global nonbacterial gastroenteritis that can be spread through contamination of food, water, fomites, or direct contact, and also via aerosolization (Fankhauser et al., 2002; Widdowson et al., 2005; Godoy et al., 2006). HuNoVs are also single-stranded RNA viruses that are small 27 to 32 nM in size that belong to the *Caliciviridae* family. However, HuNoVs are enclosed in a non-enveloped capsid, unlike SARS-CoV-2 that is enveloped. UV-C inactivation data on the HuNoV genogroups is limited due to the lack of available cultivation methods to obtain high infectious titers (Doultree et al., 1999; Ettayebi et al., 2016; Estes et al., 2019). Thus, reverse transcription quantitative polymerase chain reaction (RT-qPCR) is widely-used for assessing survivor populations of HuNoVs after treatment. However, research studies showed overestimation of survivors with RT-qPCR in comparison to virus infectivity plaque assays (Rönnqvist et al., 2014; Wang et al., 2014; Walker et al., 2019). As an alternative, cultivable animal viruses (caliciviruses, echoviruses and murine norovirus (MNV)) have been used as surrogates to determine UV-C inactivation of HuNoVs (Thurston-Enriquez et al., 2003; de Roda Husman et al., 2004; Lee et al., 2008; Park et al., 2011), but proper selection of surrogates which mimic the UV-C inactivation characteristics of HuNoVs is required to evaluate kinetics and scale up validation studies.

Furthermore, it is well known that microorganisms respond to UV exposure at rates defined in terms of UV rate constants (Patras et al., 2020). The slope of the logarithmic decay curve is defined by the rate constant, which is designated as *k*. The UV rate constant *k* has units of cm^2^/mJ or m^2^/J and is also known as the UV susceptibility. It can be also defined as D_90_ or D_10_ [dose for 90% inactivation or 10% survival] as the primary indicator of UV susceptibility. UV dose is expressed as J/m^2^ or mJ/cm^2^ (Patras et al., 2020). The varied microbial sensitivity to ultraviolet light (UV) among species of microbes, is due to several intrinsic factors including physical size, presence of chromophores or UV absorbers, presence of repair enzymes or dark/light repair mechanisms, hydrophilic surface properties, relative index of refraction, specific UV spectrum (broad band UVC/UVB vs. narrow band UVC), genome based parameters; molecular weight of nucleic acids, DNA conformation (A or B), G+C %, and % of potential pyrimidine or purine dimers (Kowalski et al., 2009).

The physical size of a virus bears no clear relationship with UV susceptibility, except that for the largest viruses, as size increases, the UV rate constant tends to decrease slightly (which is likely the result of UV scattering) (Kowalski et al., 2009). There is no thorough literature available on the above-mentioned optical parameters, hydrophilic surface properties and repair mechanisms relating to UV sensitivity. On the other hand, genome sequences of UV susceptibility can be easily retrieved from genome databases and the development of genomic models based on the above mentioned genome-based parameters is feasible to predict the UV susceptibility of ssRNA viruses, which include pathogenic novel viruses (such as SARS-CoV-2) and cultivation-challenging HuNoVs.

Our hypothesis is that predicting UV-C inactivation based on genomic modeling, will enable the determination of surrogates to be used in UV-C validation studies. In the present study, we attempted to develop a genomic model to predict and compare the UV sensitivity of enveloped SARS-CoV-2 and non-enveloped HuNoVs and to determine their suitable surrogates for use in UV-C process validation.

## 2 Materials and Methods

### 2.1 Collection of reported ssRNA viruses UV_254_ sensitivity (D_90_ values)

We collected UV sensitivity of ssRNA viruses form published studies and carefully selected D_90_ values (Table 1). The selection was based on the careful assessment of methods that were used to generate the UV dose response curves. The UV sensitivity of an ssRNA virus is determined via a dose-response curve, with the log_10_ survivors as a function of UV dose and represented as D_90_.

**Table 1.** Reported UV sensitivity (D_90_) data for ssRNA viruses

| Virus | Average $D_{90}$ (J/m<sup>2</sup>) | Reference Source |
| --- | --- | --- |
| Murine sarcoma virus | 190 | Kelloff et al., 1970; Nomura et al., 1972 |
| Bacteriophage MS2 | 183<sup>a</sup> | Malayeri et al., 2016 |
| Moloney murine leukemia virus | 115 | Nomura et al., 1972 |
| Murine norovirus (MNV-1) | 100 | Lee et al., 2008; Park et al., 2011 |
| Coxsackievirus | 79 | Battigelli et al., 1993; Gerba et al., 2002; Shin et al., 2005 |
| Human parechovirus | 75 | Gerba et al., 2002 |
| Polio virus | 73 | Gerba et al., 2002; Thompson et al., 2003; Lazaroova and Savoye, 2004; Shin et al., 2005; Simonet and Gantzer, 2006 |
| Canine calicivirus (CCV) | 67 | de Roda Husman et al., 2004 |
| Feline calicivirus (FCV) | 60 | Thurston-Enriquez et al., 2003; de Roda Husman et al., 2004; Park et al., 2011 |
| Sindbis virus | 55 | Zavadova and Libikova, 1975; von Brodrotti and Mahnel, 1982 |
| Venezuelan equine encephalitis virus | 55 | Smirnov et al., 1992 |
| Western equine encephalomyelitis virus | 54 | Dubin et al., 1975 |
| Hepatitis A virus | 51 | Wilson et al., 1992; Battigelli et al., 1993; Wiedenmann et al., 1993 |
| Semliki forest virus | 25 | Weiss and Horzinek, 1986 |
| Measles virus | 22 | Stefano et al., 1976 |
| SARS-CoV-1 | 18<sup>b</sup> | Kariwa et al., 2006 |

**Note:** Average $D_{90}$ values refers average of reference source studies; <sup>a</sup>average value of all (45) MS2 reports; <sup>b</sup>estimated value at 90 % of light transmission and initial linear kinetics

### 2.2 Determination of genomic parameters; genome size, and Pyrimidine dinucleotide frequency value (PyNNFV)

The molecular size of genomes were directly obtained from available NCBI genome database (Table 2). PyNNFV model was developed based on the frequency of each type of pyrimidine dinucleotides (TT, TC, CT and CC) which varies based on genome sequence. Pyrimidines are almost 10 times more susceptible to photoreaction (Smithyman and Hanawalt, 1969), while strand breaks, inter-strand cross links and DNA-protein cross links form with less frequency (1:1000 of the number of dimers and hydrates) (Setlow and Carrier, 1966). Three simple rules were formulated for sequence-dependent dimerization (Becker and Wang, 1989); “i) When two or more pyrimidines are neighboring to one another, photoreactions are observed at both pyrimidines, ii) Non-adjacent pyrimidines exhibit little or no photoreactivity, and iii) Purines form UV photoproducts when they are flanked at 5’ side by two or more adjacent pyrimidine residues”. Therefore, we considered 100% probability of formation of photoreaction products when PyNN are flanked by pyrimidines on both sides and 50% probability when PyNN are flanked by purine on either side. The individual PyNNs were counted by the exclusive method (each pyrimidine considered in one PyNN combination only). Research studies showed the proportion of photoreaction products in the order of TT > TC > CT > CC (Douki, 2013), thus same sequence was followed in counting individual PyNNs. Table 3 shows the method used for PyNNFV calculation in this study. A mathematical function was written to calculate PyNNFV from the potential PyNNs exist in the genome of RNA (Equation 1).

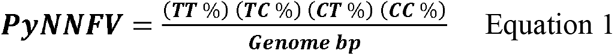

**Table 2.** Genome size and identified pyrimidine dinucleotide values for collected ssRNA viruses

| **Virus** | **NCBI ID** | **Genome (bp)** | **PyNNFV<sup>a</sup>** |
| --- | --- | --- | --- |
| Bacteriophage MS2 | NC_001417.2 | 3569 | 0.00804 |
| Murine sarcoma virus | NC_001502.1 | 5833 | 0.00807 |
| Human parechovirus | NC_001897.1 | 7348 | 0.00210 |
| Murine norovirus | NC_008311.1 | 7382 | 0.00570 |
| Coxsackievirus | KX595291.1 | 7410 | 0.00314 |
| Polio virus | NC_002058.3 | 7440 | 0.00263 |
| Hepatitis A virus | KP879217.1 | 7476 | 0.00209 |
| Feline calicivirus | NC_001481.2 | 7683 | 0.00363 |
| Moloney murine leukemia virus | NC_001501.1 | 8332 | 0.00598 |
| Canine Calicivirus | NC_004542.1 | 8513 | 0.00345 |
| Semliki forest virus | NC_003215.1 | 11442 | 0.00141 |
| Venezuelan equine encephalitis virus | NC_001449.1 | 11444 | 0.00153 |
| Western equine encephalomyelitis virus | NC_003908.1 | 11484 | 0.00151 |
| Sindbis virus | NC_001547.1 | 11703 | 0.00149 |
| Measles virus | NC_001498.1 | 15894 | 0.00134 |
| SARS-CoV-1 | NC_004718.3 | 29751 | 0.00067 |

<sup>a</sup>Pyrimidine dinucleotide frequency value**Table 3.** Calculation of PyNNFV value for SARS-CoV-2

**Table 3.** Calculation of PyNNFV value for SARS-CoV-2

| **Parameter** | **TT** | **TC** | **CT** | **CC** |
| --- | --- | --- | --- | --- |
| **PyNNs<sup>a</sup>** | 2454 | 1020 | 881 | 535 |
| **PyNNs flanked with purine<sup>a</sup>** | 773 (ATT) | 324 (ATC) | 298 (ACT) | 288 (ACC) |
|  | 412 (TTA) | 250 (TCA) | 244 (CTA) | 90 (CCA) |
|  | 530 (GTT) | 174 (GTC) | 187 (GCT) | 84 (GCC) |
|  | 230 (TTG) | 37 (TCG) | 91 (CTG) | 10 (CCG) |
| **PyNNs flanked with purine** | 1945 | 785 | 820 | 472 |
| **PyNNs flanked without purine** | 509 | 235 | 61 | 63 |
| **Probability of each PyNN<sup>b</sup>** | 1481.5 | 627.5 | 471 | 299 |
| **PyNNs %<sup>c</sup>** | 4.956341 | 2.099294 | 1.575725 | 1.000301 |
| **Genome size** | 29891 |  |  |  |
| **PyNNFV** | 0.000549 |  |  |  |

Note: a – values were counted using exclusive method (once one doublet or triplet is located in the genome, it is excluded from participating in other dimers); b – Overall probability of each PyNN is calculated by considering 50 % probability (0.5) for PyNNs flanked with purine and 100 % probability (1.0) for PyNNs flanked without purine; c – PyNNs % was determined by calculating the % of probability of PyNNs in total genome.

The PyNNFVs from complete genome sequences of 16 ssRNA viruses and corresponding reported D_90_ values were used to plot a model graph. Then, the correlation between PyNNFVs and D_90_ values was analyzed by fitting the appropriate regression model (linear regression).

## 3 Results and Discussion

Table 1 shows the median D_90_ values collected from UV-C inactivation studies of various ssRNA viruses. The data was selected from the studies conducted in transparent medium (water or phosphate buffer saline), followed standard method for UV dose calculation (Bolton and Linden, 2003). The D_90_ values reported for ssRNA viruses ranged from 18 J/m^2^ for SARS-CoV-1 to 190 J/m^2^ for murine sarcoma virus. Genomic parameters; genome size, PyNNFVs of respective viruses were shown in Table 2. The values are in the range of 3569 bp to 29751 bp for genomic size; 0.00067 to 0.00807 for PyNNFV.

### 3.1 Genomic models to predict UV-C sensitivity of ssRNA viruses

To determine the relationship between genome size and UV-C sensitivity, the D_90_ values were plotted against the genome size of various ssRNA viruses (Figure 1). The data was best fitted to log linear regression model with r^2^ = 0.63. The results revealed that decisive relationship between genome size and UV sensitivity across the range (3569 – 29751 bp).

**Figure 1.**
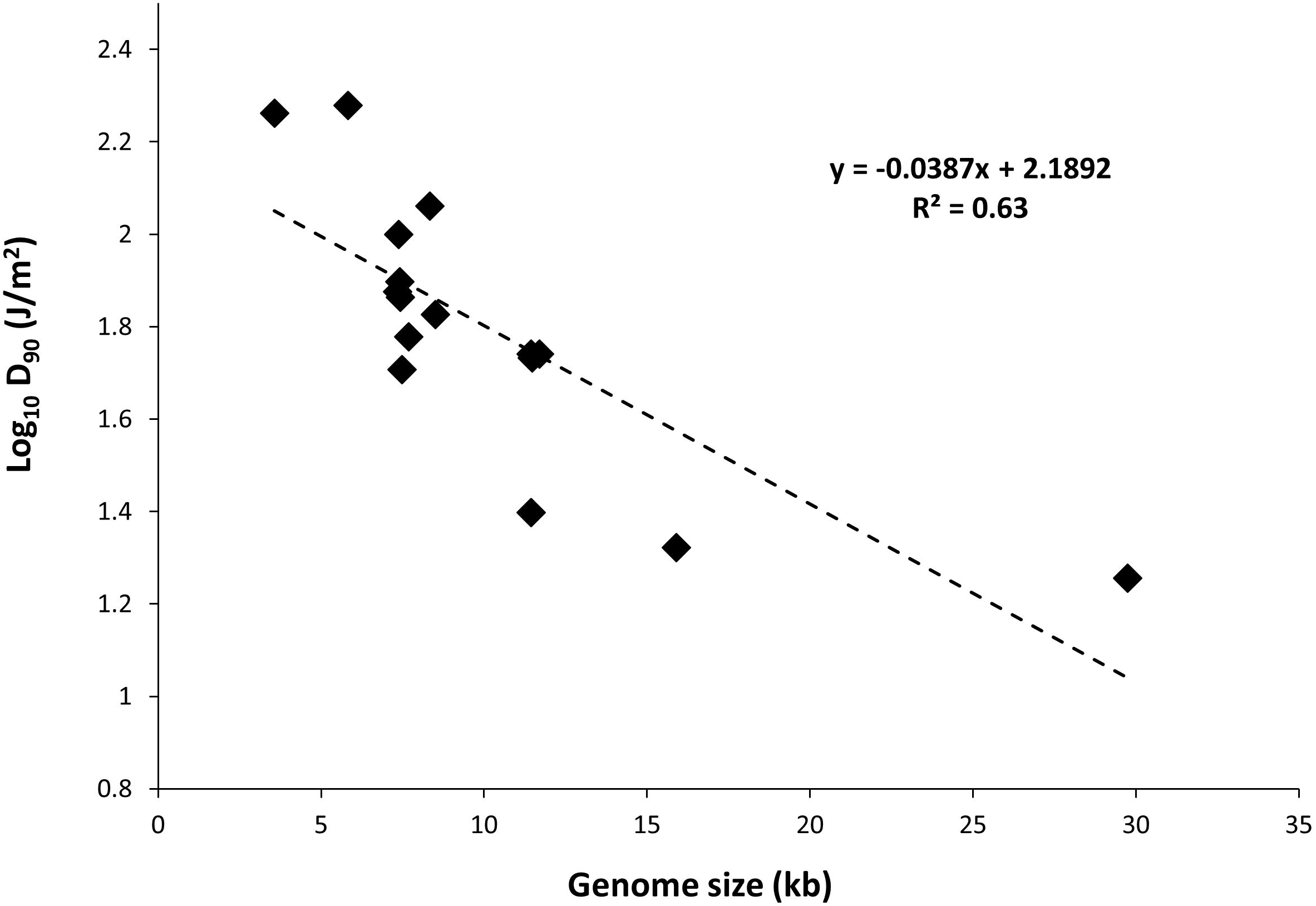
Plot of genome size versus UV-C sensitivity of ssRNA viruses

Further to evaluate the influence of base composition and sequence along with genome size on UV-C sensitivity, the D_90_ values were plotted versus pyrimidine dinucleotide frequency value (PyNNFV) (Fig 2). Linear regression model was best fitted with r^2^ = 0.90. Therefore, the results show good relationship between PyNNFV and UV-C sensitivity of virus. The following linear regression equation shows the correlation between D_90_ values and PyNNFV.

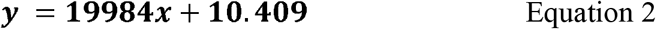

**Figure 2.**
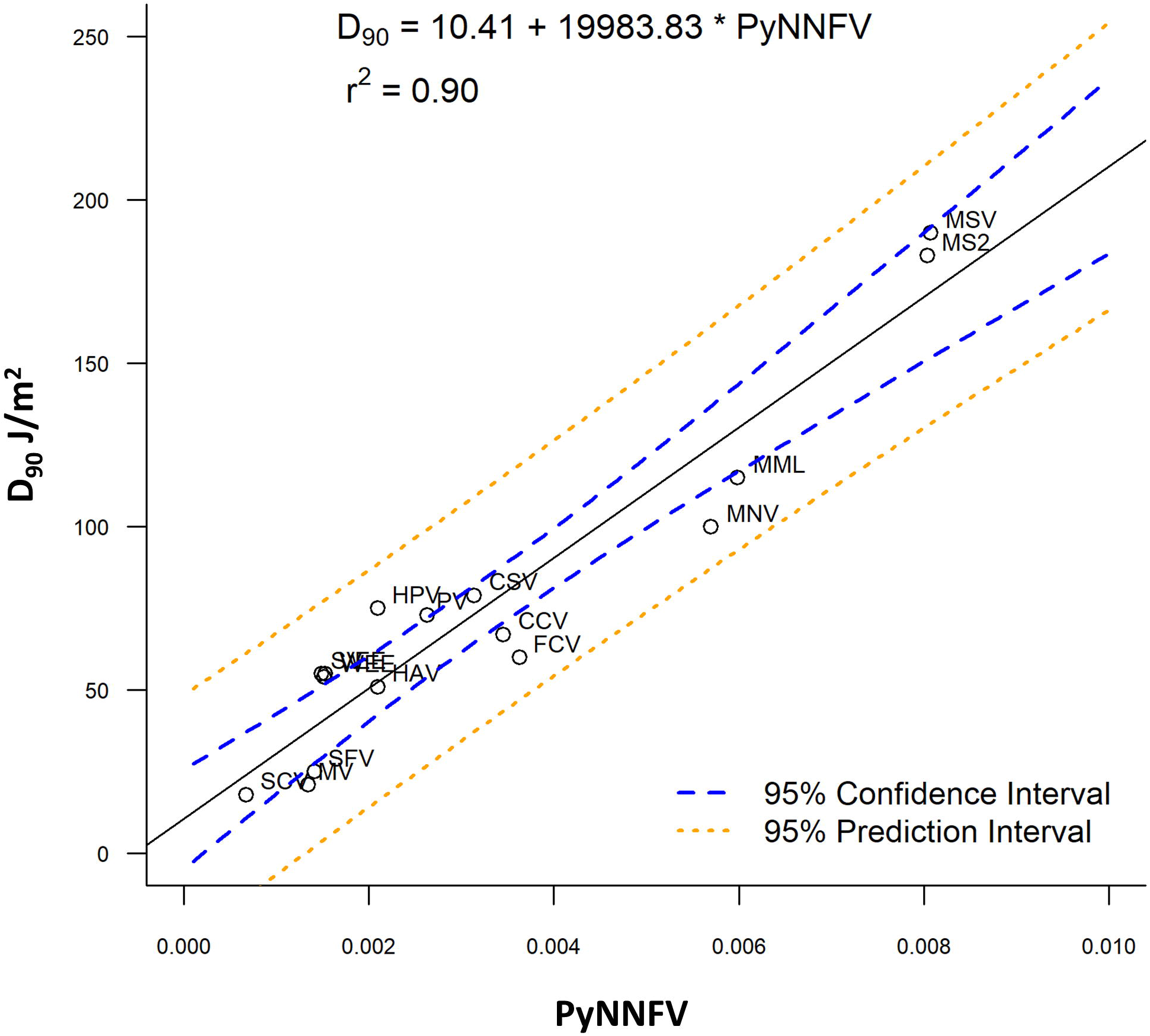
Plot of PyNNFV versus UV-C sensitivity of ssRNA viruses

Also, to predict the distribution of UV-C sensitivities and estimates of the true population mean using this model, 95% prediction and confidence intervals were shown in Figure 2. To confirm the adequacy of the fitted model, studentized residuals versus run order were tested and the residuals were observed to be scattered randomly, suggesting that the variance was constant. It can be indicated from Figure 3 that predicted values were in close agreement with the experimental values and were found to be not significantly different at p > 0.05 using a paired t-test. Despite some variations, results obtained predicted model and actual experimental values showed that the established models reliably predicted the D_90_ value. Therefore, the predictive performance of the established model can be considered acceptable. The applicability of the models was also quantitatively evaluated by comparing the bias and accuracy factors (Table 4, equation 3 & 4).

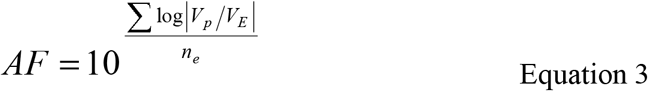

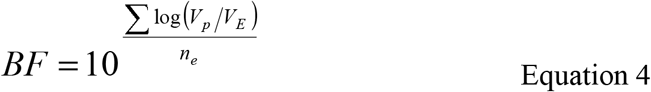

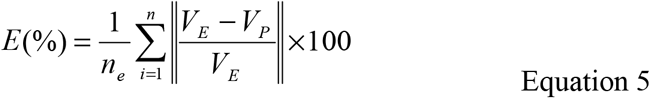

**Table 4.** Accuracy factors (AF) and Bias factors (BF) for D_90_ values in the regression analysis

| Virus | AF | BF | E (%)<sup>a</sup> |
| --- | --- | --- | --- |
| Bacteriophage MS2 | 1.02 | 1.02 | 2.18 |
| Feline Calicivirus | 0.9 | 1.11 | 12.71 |
| Coxsackievirus | 1.03 | 1.03 | 2.48 |
| Canine Calicivirus | 0.94 | 1.06 | 6.17 |
| Semliki forest virus | 0.86 | 1.16 | 18.18 |
| Murine sarcoma | 1.03 | 1.03 | 3.22 |
| Measles virus | 0.83 | 1.21 | 25.67 |
| SARS-CoV-1 | 0.91 | 1.1 | 10.88 |
| Murine Noro Virus | 0.93 | 1.08 | 8.09 |
| Moloney murine leukemia virus | 0.96 | 1.04 | 4.33 |
| Human parechovirus | 1.13 | 1.13 | 10.09 |
| Western equine encephalomyelitis virus | 1.1 | 1.1 | 8.25 |
| Venezuelan equine encephalitis virus | 1.1 | 1.1 | 8.55 |
| Sindbis virus | 1.11 | 1.11 | 9.03 |
| Hepatitis A Virus | 0.99 | 1.01 | 0.82 |
| Polio Virus | 1.05 | 1.05 | 4.62 |

<sup>a</sup>Average mean deviation**Table 5** Predicted of UV sensitivity with respect to dimerization values of target ssRNA viruses

**Figure 3.**
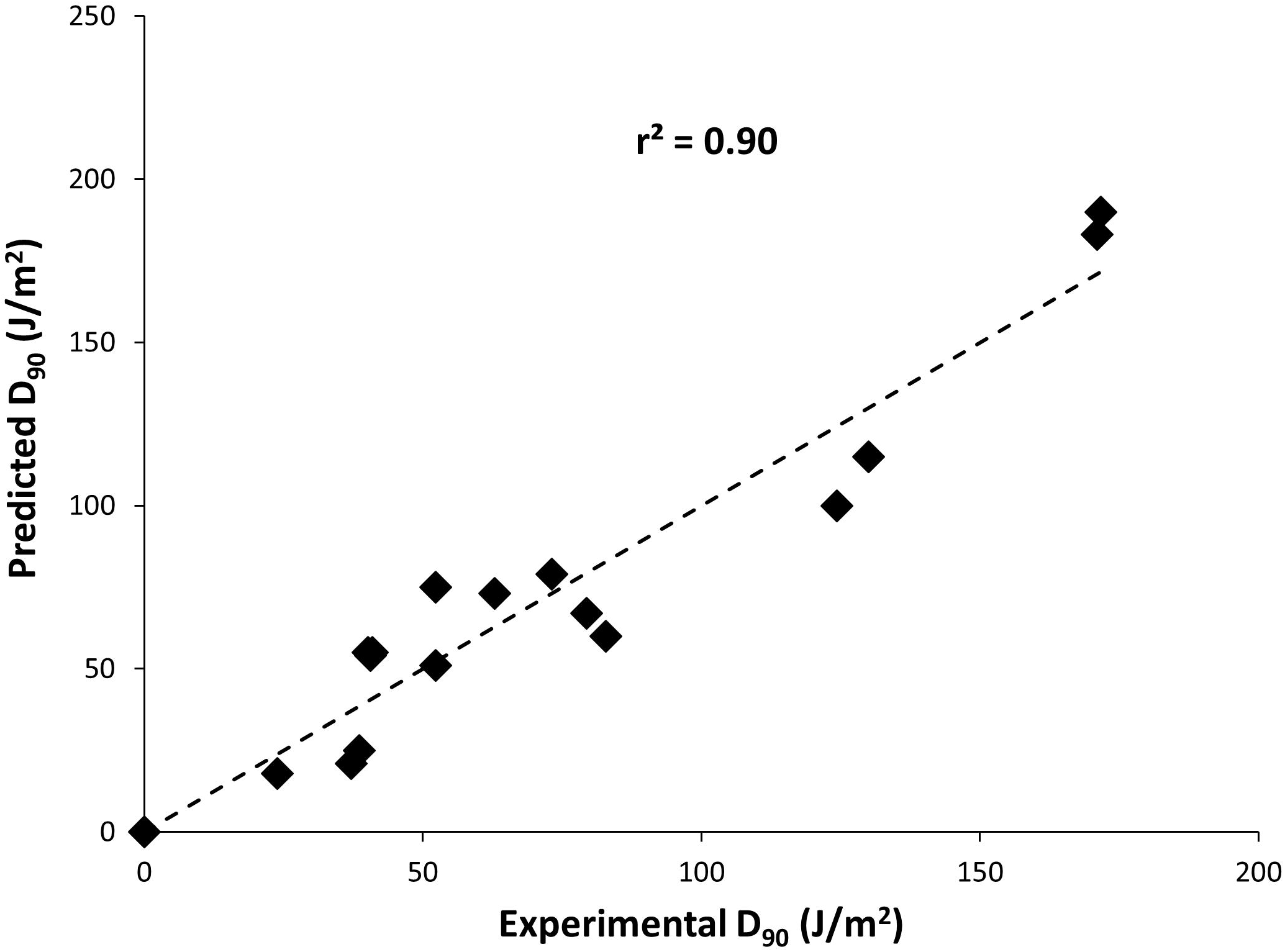
Plot of model predicted D_90_ values versus Experimental D_90_ values

The average mean deviation (*E %*) were used to determine the fitting accuracy of data (equation 5). Where, *n_e_* is the number of experimental data, *V_E_* is the experimental value and *V_P_* is the predicted value.

In most cases, as shown in Table 4, the accuracy factor (AF) values for the genomic model were close to 1.00, except for Measles virus (0.83), Semliki forest virus (0.86). The bias factor (BF) values for the predicted models were also close to 1.00, ranging from 1.02 to 1.21 for the parameter studied. These results clearly indicate that there was a good agreement between predicted and observed D_90_ values. Ross et al. (2000) stated that predictive models ideally would have an AF=BF=1.00, indicating a perfect model fit where the predicted and actual response values are equal. However, typically, the AF of a fitted model will increase by 0.10–0.15 units for each predictive variable in the model (Ross et al., 2000). Genomic model, as in this study, that forecasts a response may be expected to have AF and BF values ranging from 0.83 to 1.21 or an equivalent percentage error range of 0.82% to 25.67%.

### 3.2 Prediction of UV sensitivity of various corona viruses and human noroviruses

Owing to good model fitting, the PyNNFV genomic model was used to predict UV sensitivity of coronaviruses including SARS-CoV-2 and different HuNoV genogroups. PyNNFV values of target viruses were calculated from genomic sequences obtained from the NCBI database. The UV sensitivities were predicted by substituting PyNNFV value in equation 2. Table 5 shows PyNNFV values and corresponding predicted D_90_ values of target viruses. Predicted D_90_ of SARS-CoV-2 virus (21.4 J/m^2^) (Table 5) is closer to the estimated D_90_ of SARS-CoV-1 (18 J/m^2^) from the experimental study (Table 1). Kariwa et al. (2006) irradiated 2 mL of SARS-CoV-1 in 3-cm petri dishes without stirring UV-C light at 134 µW/cm^2^ for 15 min, and observed reduction in infectivity from 3.8 × 10^7^ to 180 TCID_50_/mL with equivalent to D_90_ value of 226 J/m^2^. In contrast, Darnell et al. (2004) showed 4 log reduction of SARS-CoV-1 at UV-C exposure of 4016 µW/cm^2^ for 6 min which is equivalent to D_90_ value of 3610 J/m^2^. The authors conducted the experiment in 24 well plate containing 2 mL virus aliquots without mixing. These two studies neither calculate the average irradiance nor provide conditions for uniform UV-C dose distribution throughout the test fluid and thereby reported higher values. The model predicted D_90_ value of MERS-CoV (28.1 J/m^2^) that is found to be higher than SARS-COV-2, whereas murine hepatitis coronavirus (MHV) strains showed similar UV-C sensitivity (D_90_ values = 20.3 to 21 J/m^2^). For α- and γ-coronaviruses, the predicted D_90_ values (17.8 to 18.3 J/m^2^) were lower than the β-coronaviruses. Saknimit et al. (1988) demonstrated the efficiency of UV-C irradiation on the inactivation of MHV and CCV coronaviruses using 15 W UV-C lamp at a distance of 1 m and reported efficient UV-C inactivation after 15 min treatment. From this data, the estimated D_90_ values for MHV and CCV (γ-coronavirus) were 17 and 15 J/m^2^, respectively, and observed to be slightly lower (~20 %) than the model predicted values (Table 5). Overall the results show that coronaviruses are highly sensitive to UV-C light than other ssRNA viruses reported in Table 1. From the UV sensitivity data obtained using the genomic model, it was observed that UV doses ranging from 90-141 J/m^2^ is required for 5 log reduction of human pathogenic coronaviruses (SARS-CoV-1, MERS-CoV, 2019-nCoV). Here we demonstrate an example of UV exposure using a low-pressure mercury lamp. If the UV-C lamp source provides an average irradiance of 0.4 mW/cm^2^ or 4 W/m^2^ (under uniform dose distribution conditions), a mere 35 second treatment is adequate to inactivate β-coronaviruses (99.999% or 5 log reduction).

**Table 5.**
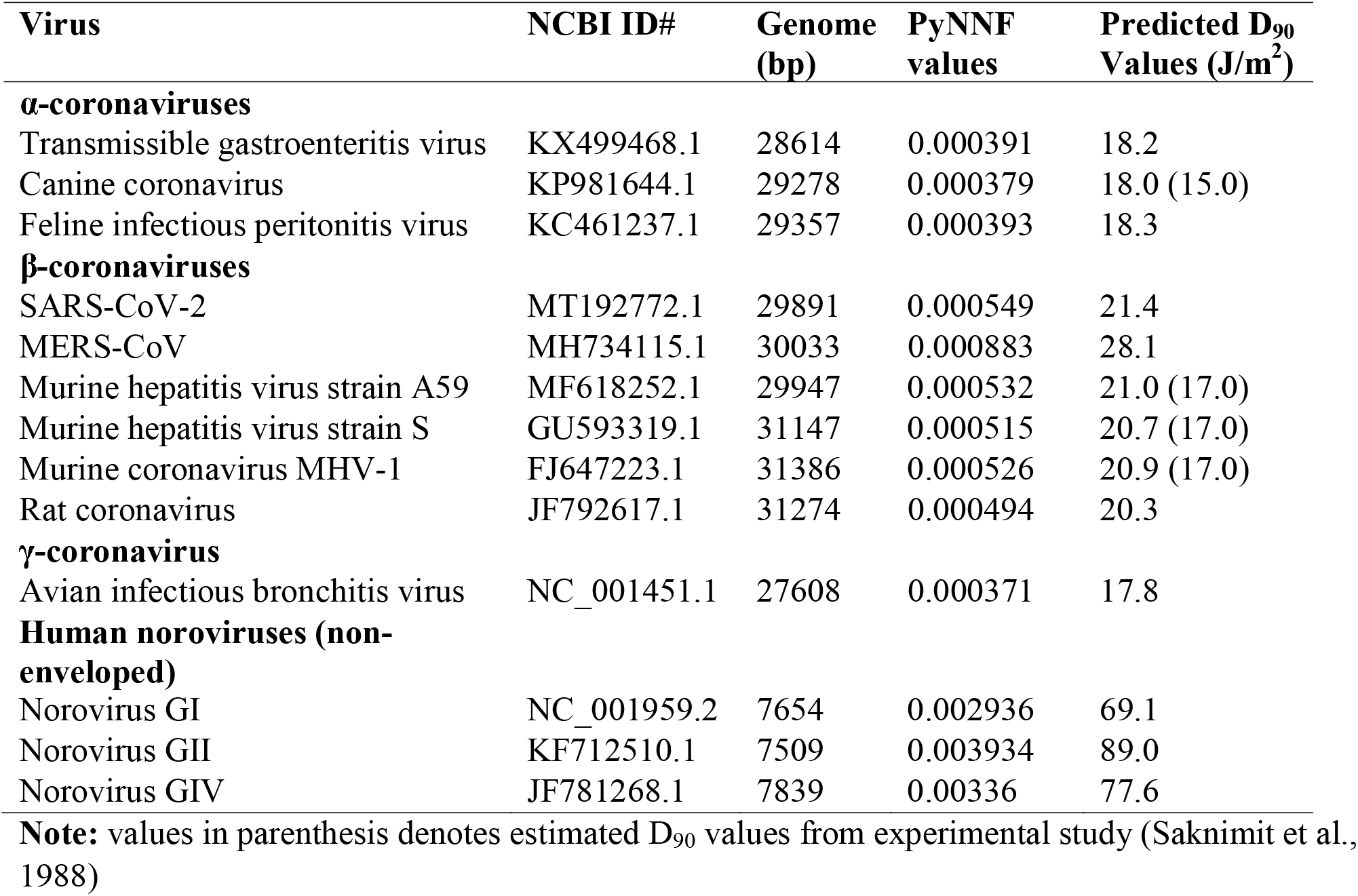
Predicted of UV sensitivity with respect to dimerization values of target ssRNA viruses

The predicted D_90_ values of HuNoVs are 69.1, 89 and 77.6 J/m^2^ for genogroups, GI, GII, GIV, respectively (Table 5). The results revealed that the UV-C sensitivity of GII was lower with higher predicted D_90_ value in comparison to GI and GIV. To the best of our knowledge, limited experimental data is currently available on UV-C sensitivity of HuNoVs. Some research studies used RT-qPCR method to estimate MNV survivors and validated with virus infectivity assay (Rönnqvist et al., 2014; Wang et al., 2014; Walker et al., 2019). The reported validation results showed that the values obtained with RT-qPCR method are overestimates to standard virus infectivity assay (Rönnqvist et al., 2014; Wang et al., 2014; Walker et al., 2019). For instance, Rönnqvist et al. (2014) reported 4-log reduction of MNV at a UV dose of 60 mJ/cm^2^ with infectivity assay, whereas just 2-log decline of MNV and HuNoV RNA levels was found at UV dose of 150 mJ/cm^2^ with RT-qPCR method. The experimental D_90_ values of conservative surrogates (MNV, echovirus and caliciviruses) obtained via viability assay are reported to be in the range of 60-100 J/m^2^ (Table 1).

### 3.3 Identification of potential surrogates for UV-C inactivation

Validation of the UV-C inactivation kinetics of specific pathogens such as SARS-CoV-2 is not possible because of the need for sophisticated biosafety level (BSL)-3 containment, and to protect the researchers, and the public from health risk in environmental settings. For HuNoV, research on reproducible cultivable systems that obtain high titers are still on-going. Hence, criteria for the selection and application of surrogates are required to ensure that the surrogates mimic the behavior of the SARS-CoV-2 or HuNoV under specific treatment conditions, while ensuring safety of personnel and also decreasing labor, cost and time. Also, surrogates are useful in process validation studies at scale up that can reduce the uncertainties linked with UV-C dose measurement.

As seen from Table 5, the model predicted D_90_ value (~21 J/m^2^) of SARS-CoV-2 was comparable to MHV strains (nonpathogenic to humans) of β-coronavirus group (~21 J/m^2^), higher than α-coronaviruses (TGEV, CCV and FIPV) and γ-coronavirus (AIBV) (~18 J/m^2^). Also, since both SARS-CoV-2 and MHV are β-coronaviruses, MHV-strain A59 may show similar behavior under various culture conditions making it a potential surrogate for SARS-CoV-2 for UV-C inactivation kinetics and validation studies.

For HuNoVs, the predicted D_90_ values of all genogroups (69-89 J/m^2^) were higher than D_90_ values of caliciviruses (60-67 J/m^2^), echoviruses (75 J/m^2^), whereas lower than MNV (100 J/m^2^) (Table 1 & 5). Use of surrogates that exhibit similar or slightly higher D_90_ values to target pathogens can avoid the risk associated with improper inactivation, hence our results indicate that MNV is the better choice (though conservative) to validate UV-C inactivation of all HuNoVs under laboratory experimental setup conditions. Furthermore, it was not surprising that the studied enveloped ssRNA viruses including SARS-CoV-1, MERS-CoV, and SARS-CoV-2 (2019-nCoV) were not as resistant to UV-C treatments compared to the non-enveloped HuNoVs and caliciviruses used in this modeling study.

In conclusion, a predictive genomic-modeling method was developed for estimating the UV sensitivity of SARS-CoV-2 and HuNoVs. Results of the model validation showed that the developed model had acceptable predictive performance, as assessed by mathematical and graphical model performance indices. We predicted the D_90_ values by conducting extensive genomic modelling. Although the parameters reported here may suffice to estimate the UV sensitivity, experimental research directed to address various knowledge gaps identified in this study is required to maximize the accuracy of predicted models. Additional parameters will be computed to the predictive model as needed, including terms for the presence of chromophores or UV absorbers and for possible UV scattering. In the future, we plan to validate this data by demonstrating experimental UV-C sensitivity (D_90_ values) of SARS-CoV-2 in containment laboratories with biosafety level 3 (BSL3) features and for HuNoVs (when suitable cultivation systems that reproducibly provide high viral titers are easily available for use in most laboratories).

## 4 Data availability statement

The genome sequences used in this study can be found in the NCBI nucleotide database. https://www.ncbi.nlm.nih.gov/nucleotide/

## 6 Author Contributions

B.P. and A.P. conceived of the presented idea. B.P. developed the theory and performed the computations. B.P., A.P., and D.D. contributed to the interpretation and discussion of the results. B.P. and A.P. wrote the manuscript.

